# Evaluation of Inducible Gene Expression Systems for Beet Cyst Nematode Infection Assays in *Arabidopsis thaliana*

**DOI:** 10.1101/2024.04.23.590774

**Authors:** Xunliang Liu, Melissa G Mitchum

**Affiliations:** Department of Plant Pathology and Institute of Plant Breeding, Genetics, and Genomics, University of Georgia

**Author notes:** Corresponding Author: Melissa G. Mitchum. Author emails: Xunliang Liu.

**Keywords:** Plant-parasitic cyst nematode, plant development, inducible gene expression, nematode infection assay, estradiol, dexamethasone

## Abstract

Cyst nematodes co-opt plant developmental programs for the establishment of a permanent feeding site called a syncytium in plant roots. In recent years, the role of plant developmental genes in syncytium formation has gained much attention. One main obstacle in studying the function of development-related genes in syncytium formation is that mutation or ectopic expression of such genes can cause pleiotropic phenotypes making it difficult to interpret nematode-related phenotypes, or in some cases, impossible to carry out infection assays due to aberrant root development. Here, we tested three commonly used inducible gene expression systems for their application in beet cyst nematode infection assays of the model plant *Arabidopsis thaliana*. We found that even a low amount of ethanol diminished nematode development, deeming the ethanol-based system unsuitable for use in cyst nematode infection assays; whereas treatment with estradiol or dexamethasone did not negatively affect cyst nematode viability. Dose and time course responses showed that in both systems, a relatively low dose of inducer (1 μM) is sufficient to induce high transgene expression within 24 hours of treatment. Transgene expression peaked at 3-5 days post induction and began to decline thereafter, providing a perfect window for inducible transgenes to interfere with syncytium establishment while minimizing any adverse effects on root development. These results indicate that both estradiol- and dexamethasone-based inducible gene expression systems are suitable for cyst nematode infection assays. The employment of such systems provides a powerful tool to investigate the function of development essential plant genes in syncytium formation.

## Introduction

Cyst nematodes (CN) within *Heterodera* and *Globodera* are among the most economically important plant-parasitic nematodes (Jones et al., 2013). They are obligate sedentary root parasites that establish a permanent feeding site called a syncytium to obtain nutrients (Sobczak et al., 1997; Holtmann et al., 2000; Sobczak and Golinowski, 2011). Once an infective second-stage juvenile (J2) locates a host root, it uses its stylet to penetrate and migrate through the root. Eventually, the J2 selects a single cell within the central vasculature as the initial syncytial cell (ISC) (Sobczak et al., 1997). A suite of effectors injected through a hollow stylet into the selected host cell reprogram it into a syncytium. Cells surrounding the ISC are gradually incorporated through cell wall dissolution and plasma membrane fusion (Grundler et al., 1998). This process is repeated with layers of cells being incorporated, forming a large metabolically active and nutrient rich multinucleate syncytium that supports the life cycle of the nematode (Sobczak et al., 1997; Holtmann et al., 2000; Sobczak and Golinowski, 2011).

While CN effectors are considered as the driving force in syncytium induction, their functions in co-opting plant development programs is essential in triggering the profound developmental and physiological changes of the host cell (Mitchum et al., 2013; Gardner et al., 2015; Siddique and Grundler, 2018; Gheysen and Mitchum, 2019; Mejias et al., 2019; Molloy et al., 2023). Several strategies have been revealed for CN effectors to manipulate plant developmental programs. First, CNs directly release molecular mimics of plant hormones, such as CLE-like peptides and cytokinins, which activate plant innate receptors and signal through corresponding signaling pathways to promote syncytium formation (Wang et al., 2005; Lu et al., 2009; Wang et al., 2011; Siddique et al., 2015; Guo et al., 2017). Second, CN effectors directly modulate hormone homeostasis in and around the syncytial cell. For example, the CN effector 19C07 that binds to the LAX3 auxin influx protein to promote its activity and increase auxin flow into the cell at the infection site (Lee et al., 2011). PIN auxin efflux transporter proteins are also laterally relocated to redirect auxin flow to facilitate syncytium expansion (Grunewald et al., 2009). Third, CN effectors directly interact with signaling components to modulate their activities. CN effector 10A07 is phosphorylated by host kinase IPK and then translocated to the nucleus to bind to IAA16, a suppressor of auxin signaling, to potentially release corresponding auxin response factors (ARFs) to activate auxin signaling (Hewezi et al., 2015). The effector 2D01 interacts with the HAESA receptor-like kinase, and may promote syncytium formation through cell wall modification (Verma et al., 2022). Furthermore, some CN effectors have been described to modulate host gene expression by interfering with regulatory machineries like histone acetylation and splicing (Verma et al., 2018; Vijayapalani et al., 2018), or through small RNA-mediated gene expression regulation (Hewezi et al., 2008; Jaubert-Possamai et al., 2019). These examples highlight the important role of host genes in CN feeding site establishment.

Therefore, dissecting how host genes are involved in syncytium formation is essential in understanding molecular mechanisms of cyst nematode parasitism, and is a vital step towards developing new strategies for genetic engineering of nematode resistant crops (McCarter, 2009; Ali et al., 2017). Despite the extensive efforts that have been made to study the function of host genes in syncytium formation, our understanding of the detailed molecular mechanisms governing syncytium formation and maintenance is still limited, especially compared to our enormous knowledge of the genetic network of plant vascular cell differentiation (Ohashi-Ito and Fukuda, 2020). One major challenge resides in the robustness of nematode effectors in inducing syncytium formation, as disruption of multiple redundant plant genes, or even parallel signaling pathways, is often required to impair cyst nematode infection. The entanglement between syncytial cell differentiation and plant development means such mutants are likely to show pleiotropic phenotypes, including poorly developed and aberrant roots, making it difficult to assess the contribution of target genes in syncytium formation. In extreme cases, mutants or transgenic lines of genes of interest could severely distort plant development or even be lethal, rendering them unsuitable for nematode infection assays.

Inducible gene expression systems, which enable conditional gene activation or suppression at desired times and within specific cell types, provide ideal solutions to overcome aforementioned obstacles. Several inducible gene expression systems have been developed and widely used in plant research communities (Aoyama and Chua, 1997; Caddick et al., 1998; Zuo et al., 2000; Zhang, 2020), of which estradiol- and ethanol-based systems have been successfully applied in root-knot nematode infection assays (Clement et al., 2009; Olmo et al., 2020). Nevertheless, it is unknown whether these systems are compatible with CN infection assays. Here, we streamlined the beet cyst nematode (BCN; *Heterodera schachtii*) infection assay in *Arabidopsis thaliana* based on the 12-well microtiter plate system described in (Baum et al., 2000), and tested three commonly used chemical-based inducible systems. Our results demonstrated that estradiol- and dexamethasone-based systems did not adversely affect BCN infection of *Arabidopsis*, and thus are compatible with this widely used model system, whereas applying ethanol significantly diminished BCN infection and thus was found to be unsuitable for use in infection assays. Adoption of these inducible gene expression systems will enable researchers to switch target genes on/off at the point of nematode infection and provide valuable tools to evaluate the role of essential developmental genes in CN parasitism.

## Result and Discussion

The 12-well microtiter plate based BCN – Arabidopsis infection assay method has been widely used since its first description by Baum et al (2000). The system offers several advantages compared to the vertical square plate-based infection assay method (Bohlmann and Wieczorek, 2015), including but not limited to: 1) each genotype/treatment is easily randomized (Figure 1A), 2) each seedling is grown in an individual well and cysts can be counted directly in the plate using a stereoscope or inverted microscope, 3) syncytium size can be measured. To further streamline this infection assay method, we developed a VBA script for randomized block design, which assigns given numbers of genotypes, treatments, or genotype-treatment combinations to an appropriate number of 12-well plates (Supplemental file 1). The script also integrates data sorting and summarizing functions to streamline downstream data analysis.

Syncytium formation during CN infection is highly intertwined with plant development programs (Siddique and Grundler, 2018). As such, many genes that could comprise syncytium formation are also likely involved in other aspects of development, and often show defective root development when mutated. For such mutants, it is critical to distinguish if the change in CN infection is due to altered root architecture or truly due to its function in syncytium formation. Several techniques were previously used to normalize nematode infection rate to total root system (Bohlmann and Wieczorek, 2015), or number of nematodes that penetrated the root system, either using high magnification light microscopy (Piya et al., 2019), or acid fuchsin staining (Bybd et al., 1983). However, with the former method, measurement of the main root does not provide an accurate measurement of the total root system available for nematode infection, and it is difficult to measure the whole root system (Bohlmann and Wieczorek, 2015), even when seedlings are grown on a vertical plate. Furthermore, this method cannot account for differences in nematode attraction and penetration (Bohlmann and Wieczorek, 2015). Although the use of the same plates for determining penetration rate is ideal (Piya et al., 2019), this approach is labor intensive, and nematodes that are fully embedded in the root are easily missed despite using a high magnification microscope.

Alternatively, nematode penetration can be easily visualized by acid fuchsin staining (Bybd et al., 1983; Grundler, 1991). To simplify the penetration evaluation, we streamlined an in-plate staining approach to evaluate nematode penetration rate (Figure 1C), which does not require removal of the plant root system from the growth medium. This method simplified the procedure and minimized disturbance to the root system. Upon conducting a penetration time course, we found that nematode penetration rate reached a maximum at two days post-inoculation and remained steady thereafter (Figure 1D), thus all penetration assays were carried out at three to five days post inoculation in our experiments.

Another approach to combat the root defects that are unfavorable to CN infection is to use an inducible gene expression system, in which genes of interest can be overexpressed or suppressed at the point of nematode infection. Several inducible gene expression systems have been developed in plants (Zhang, 2020). We reasoned that systems based on chemical inducers such as ethanol, estrogen, or dexamethasone are more suitable for the CN infection assay for ease of application, sustained effect, and minimal effect on plant growth or nematode viability compared to heat shock inducible systems, and thus were selected as our primary target for testing. Ethanol is toxic to nematodes and concentrations as low as 0.6 M (3.5% v/v) are lethal to *C. elegans* (Davis et al., 2008). Root-knot nematode (RKN) *Meloidogyne incognita* J2s in contact with 5% ethanol solution, or exposed to ethanol vapor from 5% ethanol, showed 100% immobility and 95% mortality (Silva et al., 2017). Nevertheless, the ethanol-based inducible system has been successfully employed in a RKN infection assays (Clement et al., 2009), in which the authors determined that a working concentration of 0.2% ethanol in this system did not affect RKN activity (Clement et al., 2009). However, the application of this system for CN infection assays has not been reported, thus is worth testing if the system can be used for studying CN infection. Estrogen and dexamethasone are animal derived hormones. To assess if these hormones might affect nematode activity, we first searched for estrogen and dexamethasone receptor homologues in nematodes. For this, ligand binding domain (LBD) of human estrogen receptor (hESR1, XP_016865872.1) and rat glucocorticoid receptor (RnNr3c1, NM_012576.2) were blasted (tblastn) against several nematode genomes. No significant hit was retrieved from genomes of *Heterodera* (*H*. *glycines,* taxid:51029; *H*. *schachtii*, taxid:97005; *H. avenae*, taxid:34510; *H. filipjevi*, taxid:157853), *Globodera* (*G. rostochiensis*, taxid:31243; *G. pallida*, taxid:36090; *G. tabacum*, taxid:65954; *G. ellingtoni* taxid:1517492) or *Meloidogyne* (*M. incognita,* taxid:6306*; M. javanica,* taxid:6303; *M. hapla,* taxid:6305; *M. chitwoodi,* taxid:59747) species for either hESR1-LBD or RnNr3c1-LBD. Blast against the *Caenorhabditis elegans* genome using hESR1_LBD yielded *CeNHR-49* (NM_001264305.3) with 24% identity and 46% similarity. However, a reciprocal blast of *CeNHR-49* against the human genome yielded *hHNF4G* (AB307704.1) instead of *hESR1*. These results suggest that nematode genomes do not have mammalian estrogen or glucocorticoid receptors, consistent with previous findings that *C. elegans* genome lacks NR3 type nuclear hormone receptors (NHRs), which function as classical steroid receptors (Taubert et al., 2011). Although *C. elegans* NHR-14 has been reported to bind to estrogen and activate the expression of estrogen responsive genes *vit-2* and *vit-6*, the biological effects of these interactions are not clear (Mimoto et al., 2007).

To further test if nematode penetration or development could be affected by the aforementioned chemical inducers, we tested the effect of these inducers, namely ethanol, estrogen, and dexamethasone, on nematode penetration rate and infection rate with wild-type plants. The application of an estradiol stock solution (10 mM in DMSO) to Knop’s medium resulted in visible precipitation on the medium surface (Figure S1B). Pre-dilution of the stock solution in distilled water was turbid, as indicated by optical density measured at 600 nm, due to particle precipitation (Figure S1E), whereas this was insignificant in dexamethasone diluents (Figure S1D, E). Since estradiol is stable at high temperatures, we tried to dilute the estradiol stock solution in pre-heated water (60°C). Indeed, high temperature eliminated light dissipation caused by estradiol precipitation, although we observed a slight increase after the solution was cooled to room temperature (Figure S1C, E). As such, in nematode penetration and infection assays, estradiol stock solution was first diluted to a 25x working solution in pre-heated distilled water, and 100 μl of the working solution was added to each well containing 2.5 ml Knop’s medium. For dexamethasone and ethanol, the stock solutions were pre-diluted to 25x working concentration in room temperature water, and the same amount (100 μl) was added to each well.

For estradiol and dexamethasone, no significant effect on nematode penetration or infection rates were found when the inducers were added up to final concentrations of 10 μM (Figure 2A-B, 2D-E), the maximum concentration we found in the literature for medium applications for these inducers (Aoyama and Chua, 1997; Coego et al., 2014). In addition, no significant changes were found in syncytia size following these treatments (Figure 2C, F). These results suggest that estradiol and dexamethasone do not have adverse effects on cyst nematode feeding activities and female development.

Previously, an ethanol-based gene inducible system was successfully used for RKN infection assays, without adverse effects of ethanol on RKN activity (Clement et al., 2009). However, we have previously noticed that wiping growth chambers with 70% ethanol without thorough ventilation often resulted in low cyst nematode infection rate, indicating that even exposure to low concentrations of ethanol vapor is toxic to cyst nematodes. To account for ethanol evaporation, ethanol treatment was carried out in separate plates, instead of using the randomized block design used for estradiol and dexamethasone treatments, and ethanol was added to the medium one to three days prior to nematode inoculation, to a final concentration of 0.2% (v/v) (Clement et al., 2009). When ethanol was present, nematodes still managed to penetrate into the root, although at a slightly but not significantly lower level (Figure 2G). However, further development of nematodes and syncytia was significantly inhibited (Figure 2H). Of the total of 36 plants treated with ethanol, only two adult females (Figure 2I) and three adult males (Figure 2J) were found on four plants at 14 days post-inoculation, along with a few syncytia without visible nematodes associated (Figure 2K). These results indicated that BCN viability is severely inhibited by the presence of ethanol, and strongly suggested against the use of the ethanol-based inducible system for BCN infection assays.

Next, we set out to optimize the inducer concentration and treatment length for 12-well plate grown seedlings, using the *Pro35S::XVE>>GUS* and *ProUBQ10::GVG>>4xYFP* lines. We first attempted to determine the dose and time course response of target genes using the intensity of GUS staining or YFP fluorescent signal. Both the GUS and YFP genes are highly induced by 1 μM of estradiol or dexamethasone, respectively, within 24 hours of induction. However, for both estradiol and dexamethasone, increasing inducer concentration or incubation length did not noticeably enhance the intensity of either GUS or the YFP signal (Figure S2, S3). These results suggested that GUS staining and fluorescence microscopy techniques were probably not sensitive enough to detect the differences in gene expression levels under these conditions (Figure S2, 3). Thus, qPCR was used to determine the expression profile of inducible target genes. Consistent with GUS staining and YFP fluorescence microscopy results, for both estradiol and dexamethasone, the target genes were highly induced by as low as 1 μM of the inducers, and increasing the concentration of inducers did not significantly increase the expression level of either the *GUS* or the *YFP* gene (Figure 3A, C). Nevertheless, with prolonged incubation time, expression of both *GUS* and *YFP* genes gradually increased till 3 days post induction, and then started to decline (Figure 3B, D). These results suggested that, at least for constitutive promoters like *Pro35S* and *ProUBQ10*, inducer concentration of as low as 1 μM was sufficient to induce maximum gene expression for plants grown in 12-well microtiter plates (Figure 3A, C). For both estrogen and dexamethasone induced gene expression, the expression of the target gene peaked at about 3 – 5 days post induction (Figure 3B, D). Considering that BCN penetration saturates at two days post inoculation (Figure 1E), and that BCN needs a seven hour “preparation phase” between the selection of target cell and initiation of effector secretion (Sijmons et al., 1991; Wyss, 1992), inoculation of the nematodes at one to two days post induction would ensure most nematodes are caught at the peak expression of the target gene while trying to induce syncytium formation (Figure 3B, D). Decreased gene expression 3 – 5 days post induction could be advantageous, as it would minimize any adverse effects of the transgene on root development. However, it has been reported that BCN can survive in the root for many days without successful establishment of a syncytium (Wyss, 1992), and reduced transgene expression at later stages could allow these nematodes to escape the effect of the transgene to successfully establish a feeding site (Figure 3B, D). Nevertheless, it is worth noting that data shown here only provided a baseline for inducible gene expression in the 12-well microtiter plate based system, based on the expression profile of *Pro35S::GUS* and *ProUBQ10::4xYFP* constructs. The exact gene expression profile and time point of induction would need to be adjusted based on particular use case, such as the promoter of the transgene and the goal of the particular experiment.

## Conclusion

In conclusion, we tested three commonly used inducible gene expression systems for BCN infection assays of *Arabidopsis*, and found that the ethanol-based system is not suitable for BCN infection assays due to its toxicity, whereas the steroid hormones estradiol or dexamethasone do not have adverse effects on nematode penetration or development, and are thus compatible with BCN infection assays. Recently, using the estradiol inducible system for overexpression of *MIR165a*, a microRNA specifically targeting HD-ZIP III family transcription factors, we demonstrated a major role of HD-ZIP III TFs in BCN parasitism in *Arabidopsis* (Liu and Mitchum, 2024), which would be impossible given the high level of functional redundancy of HD-ZIP III genes in vascular development, and the severe pleiotropic phenotypes of higher order HD-ZIP III mutants (Prigge et al., 2005; Carlsbecker et al., 2010). Both estradiol- and dexamethasone-based inducible gene expression systems are widely used in the plant community, along with the plentiful molecular and genetic resources that have been developed (Coego et al., 2014; Siligato et al., 2016; Schurholz et al., 2018; Machin et al., 2019; Wang et al., 2020; Shimada et al., 2022). Employment of these inducible gene expression systems in CN infection assays provides a powerful tool to investigate the function of developmental critical host genes in syncytium formation.

## Methods

### Plant and nematode materials

*Arabidopsis* ecotype Col-0 was used in this study. *Pro35S::XVE>>GUS* (Smetana et al., 2019) and *ProUBQ10::GVG>>4xYFP* (Marques-Bueno et al., 2016)(CS2106195) have been previously published. The sugar beet cyst nematode, *Heterodera schachtii*, was propagated on greenhouse-grown sugar beets (*Beta vulgaris* cv. Monohi).

### Cyst nematode infection assay

BCN eggs were isolated and hatched as previously described (Mitchum et al., 2004). After 2 - 3 days, hatched J2 were collected and surface-sterilized with 0.004% mercuric chloride, 0.004% sodium azide and 0.002% triton X-100 solution for 7 min, washed with sterilized water six times, and then resuspended in 0.1% agarose. *Arabidopsis* seeds were sterilized and grown on modified Knop’s medium with Daishin agar (Brunschwig Chemie, Amsterdam, The Netherlands) (Sijmons et al., 1991).

For phenotypic analysis, genotypes/treatments were assigned to microtiter plates following a random block design (Supplemental file 1). A single seed was placed in each well containing 2.5 ml Knop’s medium. Plates with seeds were wrapped twice with parafilm and kept in the dark at 4 °C for 2-3 days to stratify the seeds. The plates were then moved to a growth chamber set at 24 °C with a 12 h light/12 h dark light cycle.

Fourteen days after germination, each plant was inoculated with about 200 J2 suspended in 25 μl 0.1% agarose, and rewrapped twice with new parafilm. The penetration rate was counted 3 – 5 days after inoculation according to the procedure described below. Number of J4 or adult females were counted at 14 and 30 days post-inoculation, respectively. Syncytia associated with a single female were imaged at 15 days post inoculation and measured with ImageJ (Supplemental file 2). For estradiol induction, a 10 mM estradiol stock in DMSO was first diluted to working stock solutions of 25, 50, 125, and 250 μM (Figure 2A-C) in sterile water pre-warmed to 60 °C, and 100 μl of each working stock of estradiol or DMSO was added to each well (2.5 ml Knop’s media) to achieve final concentrations of 1, 2, 5, and 10 µM (25x dilution). For dexamethasone (10 mM stock in DMSO) and ethanol (200 proof), the stock solutions were diluted to 25x of final concentration (Figure 2D – F) in room temperature sterile water, and 100 μl of diluted solution was added to each well. Chemical inducers were applied 2 days prior to nematode inoculation.

### In-plate penetration assay

For penetration rate assessment, 12-well microtiter plates were processed on the same day between three to five days post-inoculation. The number of each plate was first marked to the plate side wall with a sharp object (twizzle or scissor) to avoid plate mix up in later steps. Shoots of seedlings were then removed with scissors. Plates with lids removed were placed on a floater and kept in a 100-degree water bath for 10 min to melt the solid medium. Melted medium was immediately dumped and each well was rinsed with water twice to remove medium residues. Nematodes were stained with boiling acid fuchsin (1:100 diluted) solution (0.35% acid fuchsin in 25 ml acetic acid and 75 ml distilled water) pipetted to each well in a fume hood. The plates were allowed to sit for 1-2 min, rinsed with 95% ethanol once, and then kept in 95% ethanol for counting. Nematodes will remain stained for 1 – 2 days in 95% ethanol. If nematodes are de-stained, repeat the staining procedure and count. Samples can be stored long-term in 100% ethanol to prevent nematode destaining.

### Histology staining

For GUS staining, seedlings were scooped out of the wells of 12-well plates using a spatula, and the solid medium were carefully removed. Seedlings were then treated with 80% acetone for 15 min, washed twice with GUS staining solution (100 mM PBS, pH 7.0; 1 mM potassium ferricyanide; 1 mM potassium ferrocyanide, 1 mM EDTA, 0.6% Triton X-100), and then stained with GUS staining solution containing 1 mM X-Gluc (25 mg/ml stock solution in N,N-Dimethylformamide) at 37 °C overnight. Stained seedlings were imaged with Olympus MVX10 microscope and stitched in ImageJ.

### RNA isolation and real-time PCR

For RNA isolation, *Arabidopsis* seedlings were grown in 12-well plates for 12 days, and chemical inducers were added as described before. Shoots were cut away, and roots were scooped out of the wells of 12-well plates and were placed in sterile water. After the agar medium was carefully removed, 3 – 4 roots of the same genotype/treatment were pooled together, lightly pressed between Kimwipes to remove excess water, and collected into a 1.5 ml Eppendorf tubes before freezing in liquid nitrogen. Roots were ground in 1.5 ml Eppendorf tubes with a plastic pestle with the tube immersed in liquid nitrogen to minimize RNA degradation. RNA was then isolated with the NucleoSpin RNA Plant and Fungi kit (Macherey-Nagel, Cat# 740120.5) with on-column DNA digestion. cDNA was synthesized with the PrimeScript 1st strand cDNA Synthesis Kit (Takara, Cat# 6110A) with 1000 ng of total RNA. qPCR was carried out with the PowerUp™ SYBR™ Green Master Mix (Applied Biosystems, Cat# 25742) on a Bio-rad CFX96 Maestro Machine, using Arabidopsis *ACO3* (AT2G05710) and *eIF5B1*(AT1G76810) as reference genes (Liu and Mitchum, 2024). Primer sequences are listed in supplemental file 3.

### Statistical Analysis

Statistical analyses were performed in R using Tukey’s HSD test following ANOVA analysis.

## Supporting information

supplemental_file_2_Sync_size_measurement.txt

supplemental_file_3_primers.xlsx

supplemental_file_1_Arabidopsis-Infection-Assay-template

## Acknowledgments

Special thanks to Ben Averitt, Dean Kemp, and Kurk Lance for nematode population maintenance throughout the span of this project. We thank Dr. Ari Pekka Mähönen for providing seeds.

## Author Contributions

X.L. designed and conducted all experiments, analyzed the data, and drafted the manuscript; X.L. and M.G.M. revised the manuscript.

## List of figures

**Figure 1. The 12-well microtiter plate-based beet cyst nematode infection assay.**

**A**. 14-day-old *Arabidopsis* seedings grown in a 12-well microtiter plate. Picture taken from the top of the plate. **B.** A 14-day-old *Arabidopsis* seedling grown in a 12-well microtiter plate that was removed from the well for imaging. **C.** An example of an adult female of the BCN and an associated syncytium at 14 days post-inoculation. A yellow dashed line outlines the syncytium. N, nematode. **D.** An example image showing acid fuchsin staining of nematodes in a 12-well microtiter plate at 3 days post-inoculation. Stained J2 BCNs are indicated with a yellow arrowhead. **E.** A time course of BCN penetration rate up to five days post-inoculation. Error bars represent mean ± standard error (SE). Each dot represents an individual data point. The letter above each bar represents the statistical group with Tukey’s HSD test following ANOVA analysis. The experiment was repeated twice with similar results. bar, 2 cm for (A), 5 mm for (B), 500 μm for (C), 200 μm for (D).

**Figure 2. Effect of estradiol, dexamethasone, and ethanol on BCN viability.**

**A-C.** Estradiol treatment does not affect BCN female development **(A),** penetration rate **(B),** or syncytium size **(C). D-E.** Dexamethasone treatment does not affect BCN female development **(D),** penetration rate **(E),** or syncytium size **(F). G-H.** Ethanol treatment severely suppressed BCN female development (**G**) but not nematode penetration rate (**H**). **I-J**. Examples of a syncytium (s) associated with a BCN adult female (f) (**I**), a J4 male (m) (**J**), or not associated with a visible cyst nematode (**K**). Estradiol and dexamethasone experiments were repeated twice with similar results. The ethanol experiment was performed once.

**Figure 3. Dose response and time course of estradiol and dexamethasone induced target gene expression. A-B.** Dose response **(A)** and time course **(B)** of estradiol-induced *GUS* gene expression. **C-D**. Dose response (**C**) and time course (**D**) of dexamethasone-induced *YFP* gene expression. Each bar graph represents mean ± SE. Letters on top of each bar represent statistical grouping obtained by Tukey’s HSD test following ANOVA analyses. Each experiment was repeated twice with similar results. Data from one rep was shown.

## List of e-Xtra figures

**Figure S1. Estradiol solubility test.**

**A-D,** application of estradiol and dexamethasone on Knop’s agar medium. 2.5 μl of DMSO (**A**), 2.5 μl of 10 mM estradiol stock solution in DMSO (**B**), 2.5 ul estradiol stock solution diluted to 100 μl with pre-heated (60 °C) sterile water (**C**), or 2.5 μl of dexamethasone stock solution (10 mM) diluted to 100 μl with 25 °C sterile water (**D**) was spotted onto the Knop’s agar medium. Direction application of estradiol stock solution resulted in precipitation of estradiol on the surface of the medium (**B**). Each column represents a replicate. E2, 17-beta-estradiol; Dex, dexamethasone**. E.** Light scattering test of estradiol and dexamethasone dilutions, measured with OD600. E2_RT, estradiol dilution with room-temperature (25 °C) sterile water; E2_Heat, estradiol dilution with pre-heated sterile water (60 °C); E2_Heat_RT, estradiol dilution with pre-heated water (60 °C) and then cooled to room temperature (25°C) before taking measurement; RT_Dex, dexamethasone diluted with room temperature (25°C) sterile water. Scale bar in A-D, 5 mm.

**Figure S2. GUS staining of *Pro35S::XVE>>GUS* inducible line after estradiol induction.**

Seedlings were grown in 12-well microtiter plates for 12 days before applying estradiol inducer, and were collected for GUS staining at the indicated time points. Scale bar, 5 mm.

**Figure S3. YFP fluorescent intensity of *ProUBQ10::GVG>>4xYFP* inducible line after dexamethasone induction.**

Seedlings were grown in 12-well microtiter plates for 12 days before applying dexamethasone inducer, and were fixed with 4% PFA at 4 °C overnight at the indicated time points. Images were taken with Olympus MVX10 fluorescence stereoscope. Scale bar, 5mm.

## List of supplemental files

**Supplemental file 1.** *Arabidopsis* infection assay template.

**Supplemental file 2.** A script to streamline syncytium size measurement.

**Supplemental file 3.** Primers used in this manuscript.

