## supplemental_file_2_Sync_size_measurement.txt for "Evaluation of Inducible Gene Expression Systems for Beet Cyst Nematode Infection Assays in *Arabidopsis thaliana*"

// a script for measuring syncytium size;
// Change the file extenstion to .ijm and drag it into image to use.
// Imges should be named in such a way to label the location of the syncytium, for example 'R1P01A1_1.tif', where,
//    R1 represcents Rep 1, P01 represcents Plate #1, A1 reprecents well location, and _1 to represcent different images from the same well.
//    These information can then be extracted to match to the infection data sheet produced by supplemental file 1 to get genotype/treatment info.
// at least the first image should have a horizontal or vertical scale bar in white or black color. 
//    * if no scale bar was found in an image, it will use the scal bar of the previous image. 
//    * notics that the function to find scale bar is not very robust and some time may not work. \
//      try to make the scale bar skinny and avoid end style.
//      Always check the scale bar setting under the ./bar folder to make sure scale bars are set correctly.
// Once running, it will open image one by one and promp user to outline the cell. 
//    after seletion, a .roi file will be saved to ./roi/ folder;
//    user can exit in at anytime and the program will continue with the next image when ran again;
//    after all images are processed, it will prompt user to review or continue with measuring;
//    if 'review' is selected, it will go through images and user can edit cell outlines; 
//    if 'continue' is selected, it will measure syncytium size and results will be saved to \
//       ./result folder, in a TAB seperated .txt file;
//    an image with overlayed cell outliner will be saved to ./overlayed folder;

// last modified: 2024-02-01
// Xunliang Liu
//

var bar_set, result_file, error_handle;
var bar_direction, bar_color, bar_unit, bar_length;
var path, bar_path, roi_path, result_path, overlayed_image_path;
var temp_bar_width, temp_bar_height;

getDateAndTime(year, month, dayOfWeek, dayOfMonth, hour, minute, second, msec);
today = IJ.pad(year,4) + "-" + IJ.pad(month+1,2) + "-" + IJ.pad(dayOfMonth,2) + " " + IJ.pad(hour,2) + ":" + IJ.pad(minute,2) + ":" + IJ.pad(second,2);
file_prefix = IJ.pad(year,4) + "-" + IJ.pad(month+1,2) + "-" + IJ.pad(dayOfMonth,2) + "_" + IJ.pad(hour,2) + IJ.pad(minute,2) + IJ.pad(second,2) + "_";

function init(path){
	//make subdirectories
	bar_path = path + "bars\\";
	roi_path = path + "rois\\";
	result_path = path + "Results\\";
	overlayed_image_path = path + "Overlays\\";
	if (!File.exists(bar_path)) File.makeDirectory(bar_path);
	if (!File.exists(roi_path)) File.makeDirectory(roi_path);
	if (!File.exists(result_path)) File.makeDirectory(result_path);
	if (!File.exists(overlayed_image_path)) File.makeDirectory(overlayed_image_path);
}

function setParameters(){
  Dialog.create("Scale Bar Settings");
  Dialog.addChoice("Bar Direction:", newArray("Horizontal", "Vertical"));
  Dialog.addChoice("Bar Color:", newArray("black", "white"));
  Dialog.addChoice("Bar Unit:", newArray("um", "mm", "cm", "inch", "pix"));
  Dialog.addNumber("Known Distances:", 0.5);
  Dialog.show();
  
  bar_direction = Dialog.getChoice();
  bar_color = Dialog.getChoice();
  bar_unit = Dialog.getChoice();
  bar_length = Dialog.getNumber();
}

function setupTables() {
	bar_set = bar_path + "Bar_Coordinates.txt";
	File.append("\r\n#### Bar Setting for Run: " + today, bar_set);
	File.append("Image\tCoordinates(x,y)\tWidth (pix)\tHeight(pix)\tknown distance", bar_set);
	
	//setup a table to record Measurement results
	result_file = result_path + file_prefix + "Measurements.txt";
	File.append("Image\tSynNo\tSynSize(" + bar_unit + "^2)", result_file);

	error_handle = "[Errors]";
	run("New... ", "name=" + error_handle +" type=Table");
	print(error_handle, "\\Headings:File Name\tError Message");
}

function wrapUp(check) {
	if(check == "check") return;
	//selectWindow("ROI Manager");
	//run("Close");
	selectWindow("Results");
	run("Close");
	selectWindow("Errors");
	run("Close");
}

function fileExt(filename){
	// return the extension of a file name
	s = split(filename, ".");
	if (s.length < 2) return 0;
	return s[s.length-1];
}

function baseName(filename){
	// return file name w/o extension
	ext = fileExt(filename);
	if(ext == 0) return 0;
	base_name = substring(filename, 0, lengthOf(filename) - lengthOf(ext)-1);
	return base_name;
}

function fileTypeTest(filename, ext){
	// to test if a file is 
	// 		1. with specific extension
	if (File.isDirectory(filename)) return 0;
	e = fileExt(filename);
	e = toLowerCase(e);
	ext = toLowerCase(ext);
	if(e == ext) return 1;
	return 0;
}

function ROI_Name_Parse(file_name) {
    // return an array with: label, type, id, extension;
	name_array = split(file_name, ".");
	n = name_array.length;
	if (n == 4) return name_array;
	else if (n > 4) {
		label = "";
		for(a=0; a<=n-4; a++) label = label + name_array[a] + ".";
		
		label = substring(label, 0, lengthOf(label)-1);
		type = name_array[n-3];
		id = name_array[n-2];
		ext = name_array[n-1];

		return newArray(label, type, id, ext);
	}
}

function setScale(img, _barcol, _dist, _unit){
	// set bar information for images
	// will first look for bars in dir\bars directory; if not find, will find bar in image and save an ROI in dir\bars directory;
	// bar info will be saved in table "bars".
	label = baseName(img);
	bar_file = label + ".bar.1.roi";
	if (File.exists(bar_path + bar_file)) {
		roiManager("open", bar_path + bar_file);
	} else {
		findBar(img, _barcol);
		if(selectionType() == -1) {
			File.append(img + "\tScale bar not found; Setting from previous Image is used", bar_set);
			print(error_handle, "Warning: Scale bar not found in image: " + img);
			print(error_handle, "....Scale bar setting from previous image is used");
			
			getDimensions(w, h, channels, slices, frames);
			if(bar_direction == "Horizontal") {
				makeLine(w-temp_bar_width,h,w, h);
			} else {
				makeLine(w, h-temp_bar_height, w, h);
			}
		}
		roiManager("Add");
	}
	
	roiManager("select", 0);
	Roi.getBounds(x,y,width,height);
	if(bar_direction == "Horizontal") {
		var len = width;
	} else {
		var len = height;
	}
	par = "distance="+len+" known=" + _dist + " unit=" + _unit;// + " global";
	run("Set Scale...", par);
	
    // save bar with for next image if needed;
	temp_bar_width = width;
	temp_bar_height = height;
    
    // save bar as roi file.
	saveROI("0", bar_path, baseName(img), "bar");
	run("Select None");
	roiManager("reset");
	File.append(img + "\t" + "(" + x +"," + y + ")" + "\t" + width + "\t" + height + "\t" + _dist + " (" + _unit + ")", bar_set);
}

function findBar(img, _barcol) {
	// to find bar in an image; 
	// only works if bar is in black or in white;
	// it calculates the Circ. and Roundness of particles assuming it's a rectangle;
	//    and compares the caculated results with measured Circ. and Round (deltaCirc and deltaRound);
	//    a small deltaCirc and deltaRound would be considered to be rectangle shaped thus more likely to be a bar;
	// it also compares calculated Center point of a particle to the measured Center point to make sure the particle is symmetric.
	// so far it works OK;

	run("8-bit");
	if (_barcol == "white"){ // if image has a dark background find bar in white pixels
		setThreshold(255,255,255);
	} else {			// else find bar in black pixels
		setThreshold(0,0,0);
	}

	run("Set Scale...", "distance=0 known=0 pixel=1 unit=pixel");
	run("Set Measurements...", "area centroid bounding shape limit display redirect=None decimal=3");
	run("Analyze Particles...", "size=100-Infinity pixel circularity=0.00-0.20 show=Overlay display clear in_situ");

	bar_candidates = newArray();
	for (r = 0; r < nResults; r++) {
		x = getResult("BX", r);
		y = getResult("BY", r);
		centerX = getResult("X", r);
		centerY = getResult("Y", r);
		w = getResult("Width", r);
		h = getResult("Height",r);
        if(bar_direction == "Horizontal" && h > w) continue;
        if(bar_direction == "Vertical" && h < w) continue;
		ar = getResult("AR", r);			// ar = long_axis/short_axis
		circ = getResult("Circ.", r);       // in imageJ, Circ. = 4*pi*area/(peremeter)^2
		roundness = getResult("Round", r);  // in imageJ, Round = 4*area/(pi*long_axis^2)
		solidity = getResult("Solidity", r);
		
		deltaX = abs(x+w/2-centerX);        // should be 0 if horizontally symmetric
		deltaY = abs(y+h/2-centerY);		// should be 0 if vertially symmetric
		deltaCirc = abs(3.1415926*ar/((ar+1)*(ar+1)) - circ); // the difference between a rectangle shaped Circ. to actually measured Circ. value; 
		deltaRound = abs(4/(3.1415926 * ar) - roundness);		// smaller difference means a more rectangle like shape W
		
		if (deltaCirc*deltaRound < 0.0005){ // bars with end_style will have a higher value of deltaCirc*deltaRound; 
																	  // so that this criteria is set a little high to avoid false negative;
			bar_candidates = Array.concat(bar_candidates, r+1, deltaCirc*deltaRound);     // entries meet the criteria will be added to a array;
		}
	}

	if (bar_candidates.length < 2) {                  // if no bar found, clear result window, close the image, and exit function;
		if (nResults > 0) IJ.deleteRows(0, nResults-1); // clear result window
		print("bar not found\t", img);
		return;
	} else if (bar_candidates.length > 2) {
		// if more then 2 entries meet the bar criteria;
		// find the smallest dCirc * dRound, set that one as a bar;
		rnk = Array.rankPositions(bar_candidates);
		idx = rnk[0]-1;
		bar_id = bar_candidates[idx]-1;
	} else {
		bar_id = bar_candidates[0]-1;
	}

	x1 = getResult("BX", bar_id);
	y1 = getResult("BY", bar_id);
	barw = getResult("Width", bar_id);
	barh = getResult("Height",bar_id);

	selectWindow(img);
	resetThreshold();

	if(barw > barh) {
		makeLine(x1,y1,x1+barw,y1); 
	} else {
		makeLine(x1,y1,x1,y1+barh); 
	}
	
	IJ.deleteRows(0, nResults-1); // clear result window
}

function addBar(image, length, thickness, color, position, direction, scale) {
	selectWindow(image);
	//Overlay.remove;
	getDimensions(width, height, channels, slices, frames);
	label = File.getNameWithoutExtension(image);

	//set bar parameters
    position = toLowerCase(position);
    direction = toLowerCase(direction);
    
	length_px = length*scale;
	if(thickness == 0) thickness = Math.ceil(length_px/20);
     
   if(direction == "v" || direction == "vertical") {
   	   deltaX = thickness;
   	   deltaY = length_px;
   } else {
   	  deltaX = length_px;
   	  deltaY = thickness;
   }
    
	if(position == "bottom_right") {
		x1=width-deltaX-(width-deltaX)/30;
		y1=height-deltaY-(height-deltaY)/30;
	} else if(position == "bottom_left") {
		x1=width/30;
	    y1=height-deltaY-(height-deltaY)/30;
	} else {
		//print(" bar position not known, default to bottom_right");
		addBar(image, length, thickness, color, "bottom_right", direction, scale);
		return 0;
	}

   if(direction == "v" || direction == "vertical") {
   	  x2 = x1;
   	  y2 = y1 + deltaY;
   } else {
      x2 = x1 + deltaX;
      y2 = y1;
   }

    if(x1<=0 || x2>width || y1<=0 || y2>height) {
    	showMessage("bar out of image");
    	exit(); 
    }

    //draw bar
	setColor(color);
	setLineWidth(thickness);
	Overlay.drawLine(x1, y1, x2, y2)
	Overlay.add;
	return 0;
}

function setZoom(img, z){
	// set the image scales (zoom in factor);
	selectWindow(img);
	getDimensions(width, height, channels, slices, frames);
	_par = "zoom=" + z + " x=" + width/2 + " y=" + height/2;
	setLocation(0, 0, width*z/100+300, 730);
	run("Set... ", _par);
}

function selectROI(img) {
	// prompt user to select ROIs on the open image;
	// including some flow control;
	// return values:
	//     0: set to skip;  -1: reopen previous image; -2: do nothing; or, 
	//     number of ROIs in roiManager;
	
	label = baseName(img);
	if (!fileTypeTest(img, "tif")) return -2;
	if (File.exists(roi_path + label + ".skip.1.roi")) {   
		return -2; 
	}
	if (File.exists(roi_path + label + ".syn.1.roi")) {
		return -2;
	}	

	// if ROI file is not found for the image:
	open(path + img);
	//setZoom(img,60); // set location and size of open image
	
	while (1) {
		setTool("polygon");
		waitForUser("Please outline the syncytium \r\n" + toString(i+1) + "/" + imgs.length);
		
		if (!isOpen(img)) {
			msg = "No open image.\r\nReOpen previous Image?";
			cont = getBoolean(msg, "ReOpen", "Next Image");
			if (cont == 1) return -1;
			return -2;
		}

		if (!(selectionType() == -1)) {       // if no selection on the image
			roiManager("Add");
		} else {
			if(roiManager("count") == 0) {
				msg = "No selection found on the image.\r\nDo you want to skip this image?";
				skip_image = getBoolean(msg, "Skip", "Don't Skip");
				if (skip_image == 1) return 0; 
				else continue;
			} else {
				msg = "No selection found. \r\n Add another syncytium?";
				addmore = getBoolean(msg, "Add", "Next Image");
				if (addmore==1) continue;
				else return roiManager("count");
			}
		}
		
		addmore = getBoolean("Add another syncytium?");
		if (addmore == 0) return roiManager("count");
		run("Select None");
	}
}

function setROI(path){
    imgs = listFiles(path, "tif");
	
	for (i=0;i<imgs.length;i++){
		img = imgs[i];
		nROIs = selectROI(img);
		
		if(nROIs > 0){
			saveROI("a", roi_path, baseName(img), "syn");
		} else if (nROIs == 0) { // no selections made
			saveROI("s", roi_path, baseName(img), "skip");
		} else if (nROIs == -1) {
			i = i-1;
		} else if (nROIs == -2) {
            // do nothing
        }
        
		if (isOpen(img)) close();
	}
}


function listFiles(dir, ext) {
	//list files with the given extention;
	
	result = newArray();
	files = getFileList(dir);
	for (f = 0; f < files.length; f++){
		if(fileTypeTest(files[f],ext)) result = Array.concat(result, files[f]);
	}

	return result;
}

function loadROIs(img, ROIs) {
	// find ROIs match to a image;
	img_label = baseName(img);

	result = newArray();
	for(r = 0; r < ROIs.length; r++){
		//print(ROIs[r]);
		roi_name_array = ROI_Name_Parse(ROIs[r]);       
		roi_label = roi_name_array[0];
		if(roi_label == img_label) {
            roi_type = roi_name_array[1]; 
    		result = Array.concat(result, ROIs[r]);
		}
	}

	if(result.length > 0){
        if(roi_type == "skip") {
            print("image set to skip, ROI not loaded");
        } else {
            for(r = 0; r < result.length; r++) {
                if(File.exists(roi_path + result[r])) roiManager("open", roi_path + result[r]);
            }
        }
	} else {
		print("No match found for image: " + img);
	}
	return result;
}

function setStartPoint() {

	rois = listFiles(roi_path, "roi");
	Dialog.create("Set Start Point");
  	Dialog.addChoice("Start With:", newArray("Start Over", "Image#"));
    Dialog.addNumber("Image#",1);
  	Dialog.show();
	start = Dialog.getChoice();

    if(start == "Image#") {
        s = Dialog.getNumber();
        return s-1;
	} else {
		return 0;
	}

}

function ROI_Review(path){
	// to reveiw or edit selections;
	// 1. good, NEXT
	// 2. Edited, save/replace, next
	// 3. selection change to "skip": delete syn.roi file, make .skip.roi file
	// 4. 'skip' to valid selection: add .syn.roi file
	// 5. add more syn;
	// 6. remove syn; 

	imgs = listFiles(path, "tif");
	ROIs = listFiles(roi_path,"roi");
	s = setStartPoint();

	for(i=s; i<imgs.length; i++){
		img = imgs[i];
		open(path + img);
        //setZoom(img,60);
		label = baseName(img);
		matched_rois = loadROIs(img, ROIs);

		if(matched_rois.length > 0) {
			roi_name_array = ROI_Name_Parse(matched_rois[0]);
			roi_status = roi_name_array[1];
		} else {
			roi_status = "none";
		}
		
		selectWindow(img);
		roiManager("show all");
		waitForUser("Review/Edit ROIs; \r\n (" + (i+1) + "/" + imgs.length + ")");
        
        if(!isOpen(img)) {
            reopen = getBoolean("Image is closed. Reopen it?\n", "Go Back", "ReOpen");
            if(reopen == 1) i=i-2;
            else i=i-1;
            roiManager("reset");
            continue;
        }
        
		nROIs = roiManager("count");
        
        if(nROIs == 0 && !(selectionType() == -1)) {
            addROI = getBoolean("Save selection?");
            if(addROI == 1) {
                roiManager("add");
                nROIs = roiManager("count");
            } 
        }
        
        if(roi_status == "skip") {
            if(nROIs == 0) {
                saveROI("s", roi_path, label, "skip");
                msg = today + " Reviewed: " + img;
            } else {
                File.rename(roi_path + matched_rois[0], roi_path + matched_rois[0] + ".delete");
                saveROI("a", roi_path, label, "syn");
                msg = today + " Reviewed: " + img + " |--> " + nROIs + " ROIs added (previously skipped)";
            }
        } else {
            if(nROIs == matched_rois.length) {
                saveROI("a", roi_path, label, "syn");
                msg = today + " Reviewed: " + img;
            } else {

                if(nROIs > matched_rois.length) action = "added";
                else action = "removed";
                diff = abs(nROIs - matched_rois.length);
                
                // if !nROIs == matched_rois, mark every matched_rois as "deleted"
                // and save each ROIs in ROI manager
                for(r = 0; r < matched_rois.length; r++){
                    File.rename(roi_path + matched_rois[r], roi_path + matched_rois[r] + ".delete");
                }
                
                if(nROIs == 0){
                    saveROI("s", roi_path, label, "skip");
                    msg = today + " Reviewed: " + img + "  |--> " + diff + " ROIs" + action + " Image set to skip";
                } else {
                    saveROI("a", roi_path, label, "syn");
                    msg = today + " Reviewed: " + img + "  |--> " + diff + " ROIs" + action;
                }
            }
        }

        selectWindow(img);
        run("Close");

		
	}

}

function saveROI(idx, path, name, type){
	// idx: index of an ROI to save; 'a', save all ROIs; 's' skip, make a line and save it. 
	// will reset roiManager at the end;
	if (idx == "a"){ 
		nROIs = roiManager("count");
		for (roi_idx = 0; roi_idx < nROIs; roi_idx++) {
			seriesNo = roi_idx + 1;
			label = name + "." + type + "." + seriesNo;
			roiManager("Select",roi_idx);
			roiManager("Rename", label);
			saveAs("Selection", path + label + ".roi");
		}
	} else if(idx == "s") {
		seriesNo = 1;
		label = name + "." + type + "." + seriesNo;
		roiManager("reset");
		makeLine(0,0,200,200);
		roiManager("Add");
		roiManager("Select",0);
		roiManager("Rename", label);
		saveAs("Selection", path + label + ".roi");
	} else {
		idx = parseInt(idx);
		seriesNo = idx + 1;
		label = name + "." + type + "." + seriesNo ;
		roiManager("Select",idx);
		roiManager("Rename", label);
		saveAs("Selection", path + label + ".roi");
	}
	roiManager("reset");
}


function measureTheSize(_roi){
	name_array = ROI_Name_Parse(_roi);
	label = name_array[0];
	roi_type = name_array[1];
	roi_id = name_array[2];
	img = label + ".tif";
	img2 = label + "..tif";
	if(roi_type == "skip") {
		return;
	}

	if(File.exists(path + img)) {
		open(path + img);
	}
	else if(File.exists(path + img2)) {
		open(path + img2); //for some files, there're two dots before the file extension. 
	}
	else {
		return;
	}
	img_w = getWidth();
	img_h = getHeight();
	setScale(img, bar_color, bar_length, bar_unit);
	roiManager("reset");
	roiManager("open", roi_path + _roi);
	roiManager("select",0);

	run("Set Measurements...", "area bounding display redirect=None decimal=3");
	run("Measure");
	syn_size = getResult("Area", 0);
	File.append(label + "\t" + roi_id + "\t" + syn_size, result_file);
	IJ.deleteRows(0, nResults);

    //generate overlay image
	bar_file = name_array[0] + ".bar.1.roi";
	if (File.exists(bar_path + bar_file)) roiManager("open", bar_path + bar_file); // load bar roi
	nrois = roiManager("count");
	roiManager("select", nrois-1);
	Roi.getBounds(bar_x, bar_y, barw_px,barh_px);
	bar_xx = (img_w - barw_px) - (img_w - barh_px)/30;
	bar_yy = (img_h - 10) - (img_h - 10)/30;
    if(bar_direction == "Horizontal") {
        bar_scale = barw_px/bar_length;
    } else {
        bar_scale = barh_px/bar_length;
    }
    
    addBar(img, bar_length, 10, "red", "bottom_right", bar_direction, bar_scale);

	
	for (rs = 0; rs < (nrois-1); rs++){
		roiManager("select", rs);
		Overlay.addSelection("yellow", 2);

	}
    
	//waitForUser("test: check overlay image");
	Overlay.flatten;
	//waitForUser("test: check overlay image");
	par = "width=" + img_w/2 + " height=" + img_h/2 + " constrain average interpolation=Bilinear";
	run("Size...", par);
	saveAs("Jpeg", overlayed_image_path + baseName(_roi) + ".flatten.jpeg");
	run("Close All");
	roiManager("reset");
	}

// main code
macro SynSize {
	
	path = getDirectory("Choose a Directory");
	//print(path);
	init(path);
	setROI(path);
	
	msg = "ROIs setup done. \r\n     Press 'Continue' to measure;\r\n     Press 'Review' to review/edit selections;\r\n     Press 'Cancel' to exit";
	toMeasure = getBoolean(msg, "Continue", "Review");
	//print(toMeasure)
	if(toMeasure == 0) ROI_Review(path);
	
	setBatchMode(true);
	setParameters();
	setupTables();

	synROIs = listFiles(roi_path, "roi");
	for (i=0; i<synROIs.length; i++){
		measureTheSize(synROIs[i]);
	}
	
	wrapUp("exit");
    showMessage("Measurement done!");
	setBatchMode(false);
} 
